# How muscle ageing affects rapid goal-directed movement: mechanistic insights from a simple model

**DOI:** 10.1101/2025.07.03.663079

**Authors:** Delyle T. Polet, Christopher T. Richards

## Abstract

As humans and other animals age, passive and active muscle properties change markedly, with reduced peak tension, peak strain rate, activation and deactivation rate, and increased parallel stiffness. It is thought that these alterations modify locomotor performance, but establishing causal links is difficult when many parameters vary at once. We developed a simplified model of a single joint with two antagonistic Hill-type muscles, and varied the associated muscle parameters combinatorially over a large range. For a given parameter combination, we found optimal joint movements that minimized cumulative squared error to a target while starting and ending at rest. Emergent behaviour from the optimisations compared well to ballistic point-to-point arm movements in humans. Age-associated reductions of maximum isometric force, maximum strain rate and activation rate all had detrimental effects on performance, independent of other parameters. In contrast, deactivation time and passive parallel stiffness had no effect on performance on their own, but pronounced interactive effects with each other. Increasing stiffness reduced joint movement time at fast deactivation rates, but increased movement time at slow deactivation rates. This occurs because antagonist muscles resist the passive tension at rest, but are stretched eccentrically by the agonist, amplifying their active resistive force. Fast-deactivating muscles can avoid this resistive effect, allowing the passive stiffness to amplify accelerating force and enhance performance. In all cases, coactivation emerged during and after the braking period, and during the acceleration phase when stiffness increased. As deactivation time increased, so too did coactivation levels– but coactivation was not generally associated with a reduction in performance. Our simulations offer evidence that age-related changes in muscle strength, activation time and maximum contraction velocity could lead to reductions in ballistic performance in a goal-directed task, but the effects of increased muscle stiffness and deactivation time depend on their relative values.

## 2 Introduction

Muscle is the dominant engine of animal movement, and physiologists have long been interested in how variation in muscle contractile properties is linked to functional outcomes (James et al., 1995; Syme, 2005; Richards and Clemente, 2013; Polet and Labonte, 2024). Physiological changes in muscle are especially pronounced (and best studied) during ageing. As mammalian muscles age, they generally exhibit reductions in isometric tensile strength (Brooks and Faulkner, 1988; Brown et al., 1999; Clark and Taylor, 2011; Valour et al., 2003; Holt et al., 2016; Moran et al., 2006), slower activation and deactivation (Brooks and Faulkner, 1988; Harridge et al., 1997; Danos et al., 2016; Kiriaev et al., 2021; Moran et al., 2006; Hill et al., 2018), reductions in the effective (muscle-level) maximum strain rate (Valour et al., 2003; Raj et al., 2010; Holt et al., 2016), and increases in passive parallel stiffness (Brown et al., 1999; Wood et al., 2014; Noonan et al., 2020; Moran et al., 2006).

In addition to changes in muscle contractile properties, ageing is also associated with changes in locomotor behaviour and performance. Elderly humans tend to reach more slowly and with more effort (Welsh et al., 2007; Summerside et al., 2024), with less precision (Welsh et al., 2007), and with higher levels of cocontraction (Hortobágyi and DeVita, 2006; Seidler-Dobrin et al., 1998). Performance reductions in reaching tasks are associated with negative health outcomes, particularly fall risk (Maki and McIlroy, 2006). Establishing links between physiological changes in muscle and performance outcomes provides a theoretical basis for future research towards targeted treatment to maintain locomotor performance throughout life. However, direct links are difficult to establish, as ageing can involve multifactorial musculoskeletal and neurological changes (Tieland et al., 2018) that are difficult to fully control for in empirical studies. Which changes to muscular parameters actually lead to performance differences in everyday movement? Which are most important, and do they interact?

To address the above questions, dynamic biomechanical models can be used to explore the effects of varying parameters in a well-defined control task. Applying optimal control to these models yields behavioral predictions of motor control, providing an upper bound on performance for a given set of parameters. (Falisse et al., 2019; Polet and Bertram, 2019; Polet and Labonte, 2024). Despite their potential predictive power (Falisse et al., 2019, 2020; De Groote and Falisse, 2021), highly complex models can be difficult to analyse in detail (De Groote and Falisse, 2021; Halilaj et al., 2018), as the number of interdependent parameters becomes intractably large. In contrast, simplified models can search a large number of parameter combinations to unravel links between underlying parameters, their interactions, and their consequences for optimal behaviour (Wong et al., 2016; Polet, 2021; Koelewijn and Van Den Bogert, 2022). They also allow us to better understand the mechanisms between muscle change and performance outcomes (Murtola and Richards, 2023). As single-joint and multi-joint arm movements appear to use similar control strategies (Flash and Hogan, 1985; Kaminski and Gentile, 1989; Berret et al., 2024), we focus on a simplified point-to-point reaching movement about the elbow, allowing us to better understand the mechanisms underlying performance changes.

To that end, we propose a simplified two-muscle Hill-type model of a single joint. We assign parameters based on the human forearm and use optimal control to minimise time-integrated squared position error in a point-to-point movement. This metric of performance encapsulates both the time to reach the target, and rapid stabilisation around it. Using a large parameter sweep of isometric force, maximum strain rate, parallel passive stiffness, and activation and deactivation rates, we determine best-case performance outcomes for thousands of parameter combinations. The results of this investigation tell us which age-related changes might be associated with performance deficits, how and when they interact, and uncover causal links between muscle physiology and performance outcomes.

## 3 Methods

### 3.1 Model design

We simulated a motor control task to extend a rotational joint from one point to another, starting and ending at rest, while minimizing a performance cost within a set time. The cost is given as the time-integrated squared error from the target (endpoint) position; more time spent away from the target accrues higher cost, and so high performance (low cost) indicates a solution that can rapidly reach and come to rest at the target.

The arm, muscle and activation models are based on Murtola and Richards (2022, 2023), but considering only single degree-of-freedom flexion and extension about a frictionless elbow joint. This further simplification enables a far more thorough sweep of parameters than typically performed in musculoskeletal modelling studies. In order to keep the model simple and conceptual, the flexor and extensor muscles are identical in size, strength, muscle properties, and moment arms. The load they move is a lumped inertia equivalent to a combination of modeled forearm and hand inertias, with a fixed wrist. The motion is in a horizontal plane, such that there are no external forces. At 90^°^ flexion, both flexor and extensor are at optimal (resting) length, and the forearm is perpendicular to the upper arm (Figure 1).

**Figure 1:**
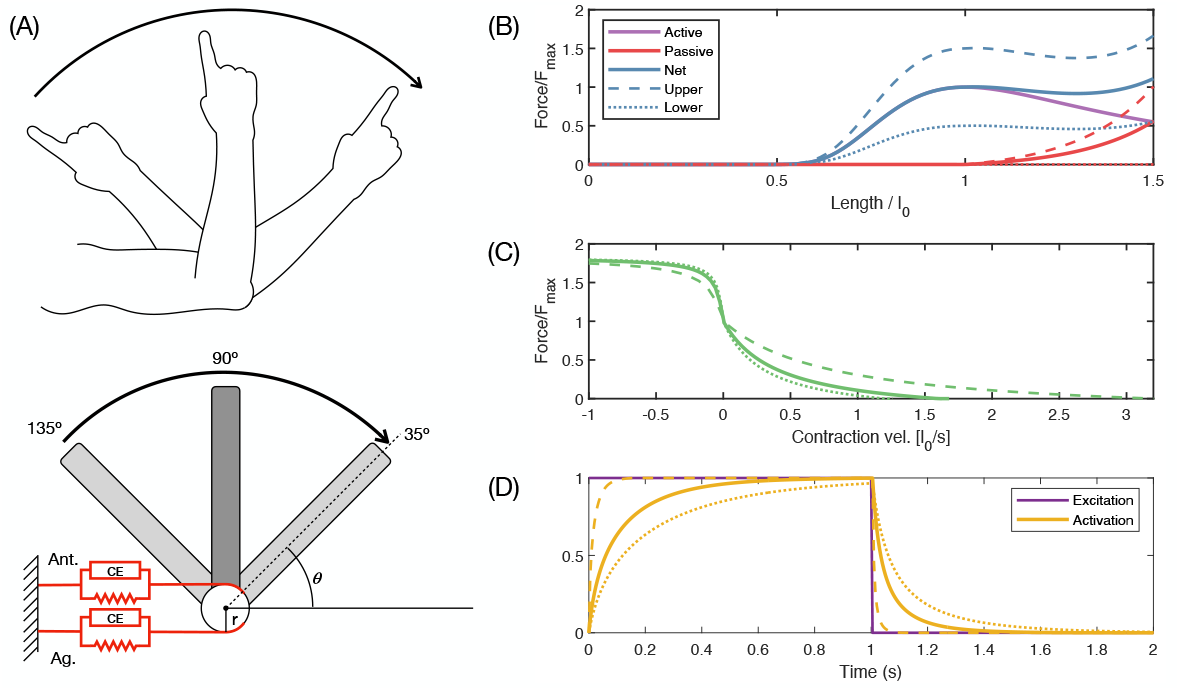
(a) Single joint model. Muscles are Hill-type with constant moment arm *r*. (b-d) Hill-type muscle and activation properties. Baseline case is indicated as a solid line, while the lower and upper ranges explored in the parameter sweep are shown as dotted lines. Parameters are specified in Tables 1 and 2. (b) Hill-type force length properties. Active force length is based on Otten (1987) and parallel passive force length based on Murtola and Richards (2023). (c) Hill-type force-velocity properties based on Otten (1987). (d) Activation model based on Thelen (2003). Mathematical representations are smoothed to remove discontinuities for trajectory optimisation; for details, see supplemental text.

#### 3.1.1 Parameter range selection

Baseline parameter values were taken from Murtola and Richards (2023). Parameter ranges to explore were based on putative changes with age observed in empirical studies. For peak tetanic force, Brown et al. (1999) found at most a 2-fold reduction for healthy older mice compared to young controls (in the Plantaris muscle), and a 5-fold reduction for “sendentary” (hindlimb offloaded) mice. We therefore considered a 3-fold range in peak tension (0.5 to 1.5 times the baseline value). The same study found at most a 2.5-fold increase in normalised passive parallel stiffness for old mice compared to young, and at most a 5.5 fold increase in hindlimb-offloaded old mice compared to young. We chose a maximum stiffness 1.8 times the baseline value (higher values led to difficulties in simulation convergence) and also considered the case of no muscle parallel stiffness.

Raj et al. (2010) compiled empirically measured changes in muscle *v*_max_ values in humans, finding they are reduced by at most 1.6 times in elderly subjects compared to young. However, an empirical study of human elbow flexion (Valour et al., 2003) reports *v*_max_ values twice as large as used by Murtola and Richards (2023). To account for this uncertainty in *v*_max_ variation, we chose a range from 0.75 to 2 times the baseline value (a 2.7-fold change).

Doherty et al. (1993) reviewed studies on muscle contractile properties with age, finding that the largest reported change in time to peak tension in older populations was at most 1.5 times the younger value in the extensor digitorum brevis (EDB), and about 1.25 times the younger value in triceps surae (TS). Time to half relaxation increased 2 times in the EDB and up to 1.2 times in the TS. However, it is challenging to map twitch-based activation parameters to a first-order model that does not reproduce twitches (Murtola and Richards, 2022), leading to uncertainty as to the most representative numerical value. Moreover, we noted parameter interactions with deactivation that warranted further investigation. Therefore, we chose a much larger variation in activation and deactivation time (from 0.15 to 2 times the baseline value) to explore these issues, while acknowledging this is much larger net change than could be reasonably expected during human ageing.

In addition to the large parameter sweep, which varied all parameters over their full ranges outlined above, we also performed a finer-grid parameter sweep, which varied only deactivation time and stiffness constants. These were varied over the same range as in the larger parameter sweep, but in 0.6 ms and 0.005 unit increments, respectively. All other parameter values were held at baseline.

### 3.2 Trajectory optimisation

For the current study, the optimisation problem is to start at a given position at rest, and choose time-varying excitation values that minimize the sum of squared position error (SSE) to a target, while satisfying dynamic constraints, and finally ending at rest. SSE was defined as

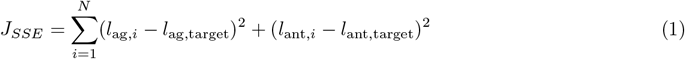

where *l*_ag,*i*_ and *l*_ant,*i*_ is the muscle length of the agonist and antagonist, respectively, at timestep *i*. For comparison to empirical EMG data, start and target positions of 120^°^ and 60^°^ respectively were used. For parameter sweeps, start and target positions of 135^°^ and 35^°^ were used. As the end position was not constrained, the objective was augmented with an additional term penalizing an end position far from the target

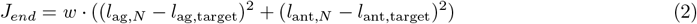

where the weighting *w* = 0.01 and *N* corresponding to the final time step. The final objective was a combination of terms, *J* = *J*_*SSE*_ + *J*_*end*_. The time horizon *T* was set as 0.4 s, chosen such that the worst-performing case could reach the target.

Force-length and force-velocity relationships are Hill-type following Otten (1987). These were smoothed where cusps and discontinuities existed (see Supplemental Information), so that exact derivatives could be calculated using CasADi (Andersson et al., 2019). Excitation-activation coupling was set using a first-order model following Thelen (2003). Muscle excitations were set as piecewise constant controls. Modeling constants and baseline parameters (Tables 1 and 2) are chosen to match Murtola and Richards (2022), with the exception of *b*_1_ in the force-length relationship. This was given the value of 2 so that the Hessian of force to length was finite everywhere. The maximum eccentric normalised force (parameter *d*_2_) was scaled to changes in *F*_max_ in order to maintain absolute eccentric force 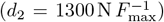, in accordance with evidence that eccentric strength remains relatively well preserved with advancing age in humans (Roig et al., 2010).

**Table 1:** Modelling constants. *For variable values, see Table 2

| Parameter | Symbol | Value |
| --- | --- | --- |
| Muscle moment arm | $r$ | 0.04 m |
| Muscle resting length | $l_0$ | 0.32 m |
| Simulation duration | $T$ | 0.4 s |
| Arm resting angle | $\theta_0$ | $90^\circ$ |
| Arm initial and final position | $[\theta_i, \theta_f]$ | $[45^\circ, 135^\circ]$ |
| Arm moment of inertia about elbow | $I$ | $6.88 \times 10^{-2} \text{ kg m}^2$ |
| Force-length shape parameters | $\mathbf{b}$ | $[2, -1.3, 0.53]$ |
| Passive parallel stiffness parameters | $\mathbf{c}$ | $[c_1, 5, c_3]^*$ |
| Slack length per $l_0$ | $c_3$ | 1.0 |
| Force-velocity shape parameters | $\mathbf{d}$ | $[4, 1.4, 30.24]$ |
| Smoothing parameter | $s$ | 500 |

**Table 2:** Baseline parameters and values tested during parameter sweeps. For the finer-grid parameter sweep, deactivation time and stiffness values were varied over the same range below, in 0.6 ms and 0.005 unit increments, respectively, while all other values were held at baseline.

| Parameter | Symbol | Baseline value | Range tested (× baseline value) |
| --- | --- | --- | --- |
| Peak active force | $F_{\max}$ | 1300 N | $[0.5, 0.8, 1.0, 1.5]$ |
| Maximum strain rate | $v_{\max}$ | $1.6 l_0 \text{ s}^{-1}$ | $[0.75, 1.0, 1.5, 2.0]$ |
| Activation time constant | $\alpha_a$ | 94.5 ms | $[0.15, 0.25, 0.5, 0.75, 1.0, 1.5, 2.0]$ |
| Deactivation time constant | $\alpha_d$ | 65 ms | $[0.15, 0.25, 0.5, 0.75, 1.0, 1.5, 2.0]$ |
| Stiffness constant | $c_1$ | 0.05 | $[0, 0.6, 1, 1.4, 1.8]$ |

The optimisation problem was discretised into 5 ms timesteps, and transcribed with the CasADi opti stack using multiple shooting with 4th-order Runge-Kutta trapezoidal integration. The trajectory optimisation problem was solved using IPOPT (Wächter and Biegler, 2006) with the MA57 linear solver. For every case in the large parameter sweep, and for the baseline case, the initial guess consisted of a linear interpolation of muscle lengths from the starting value at *t* = 0 to the target value at *t* = *T*, and zero values for all other states and controls. For the finer-grid parameter sweep, the baseline case solution was used as the initial guess (since other parameters were held at baseline values).

The optimisation proceeded for every parameter combination outlined in Table 2, a total of 3920 trials, as well as for the 399 trials of the finer-grid sweep, all of which converged to unique optimal solutions. The performance of all solutions was evaluated as the Root Mean Square Error, defined as

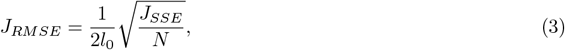

where the value of 2 allows *J*_*RMSE*_ to represent the normalised error of one muscle (as both muscles are symmetric and contribute equally to *J*_*SSE*_).

### 3.3 Regression

In order to determine which of the parameters, or their interactions, contribute to changes in performance in this dataset, we performed linear regression of the *J*_*RMSE*_ against the five muscle parameters and their interactions. A common method for determining the relative importance of individual terms in a regression model is “standardized betas” (Mizumoto, 2023), where predictors are standardized by subtracting their means and dividing by standard deviation. The regression coefficients for each term thus represent the change in the response variable from a unit (standard deviation) change in the predictor, and thus can be seen as the relative strength of the predictor in affecting the response (Mizumoto, 2023).

However, the standardized beta method assumes a Gaussian distribution of the data, whereas the predictors in this case are not randomly distributed. Instead, we normalise each muscle parameter of interest and the response variable (*J*_*RMSE*_) by subtracting its minimum value and dividing by its range. Thus, the regression coefficient for each term represents the fractional change in *J*_*RMSE*_ expected by changing the predictor from its minimum to maximum value (in the absence of interactions).

A criticism of standardized betas is that results can be misleading if predictors are strongly correlated with each other (Mizumoto, 2023). However, the predictors in this deterministic model are uncorrelated by design, and so regression coefficients should be a strong indicator of importance. Nevertheless, if different predictor ranges were used (e.g. doubling the range of *F*_max_ in the parameter sweep), then different regression coefficients could emerge, leading to different measures of importance. This issue is beyond the scope of the current study, but a critical consideration when interpreting results.

Coactivation at a given time *t* was defined as the minimum activation between the antagonist and agonist at a given timepoint (*a*_*c*_(*t*) = min(*a*_ant_(*t*), *a*_ag_(*t*))), while the average of coactivation was taken over the duration of the simulation,

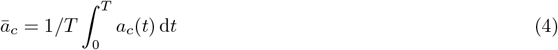

## 4 Results

Using baseline parameters (Table 1), predictive simulation of optimal elbow extension exhibits qualitative similarities to empirical ballistic data from Forgaard et al. (2013, Figure 2). In both, the elbow extension profile is smooth and settles at the target after overshooting. The simulation predicts two main features that also appear in experimental data: (a) a triphasic pattern of excitation, where agonist and antagonist excitation are out of phase in an agonist-antagonist-agonist pattern (Leib et al., 2020; Correia et al., 2022), although the simulated excitation and empirical EMG are offset in time (Figure 2); and (b) an additional final burst of antagonist excitation following the completion of the movement, which corresponds to a sustained level of excitation in the agonist and antagonist. Rapid single-joint arm movements are associated with relatively high levels of tonic coactivation following movement (Suzuki et al., 2001).

**Figure 2:**
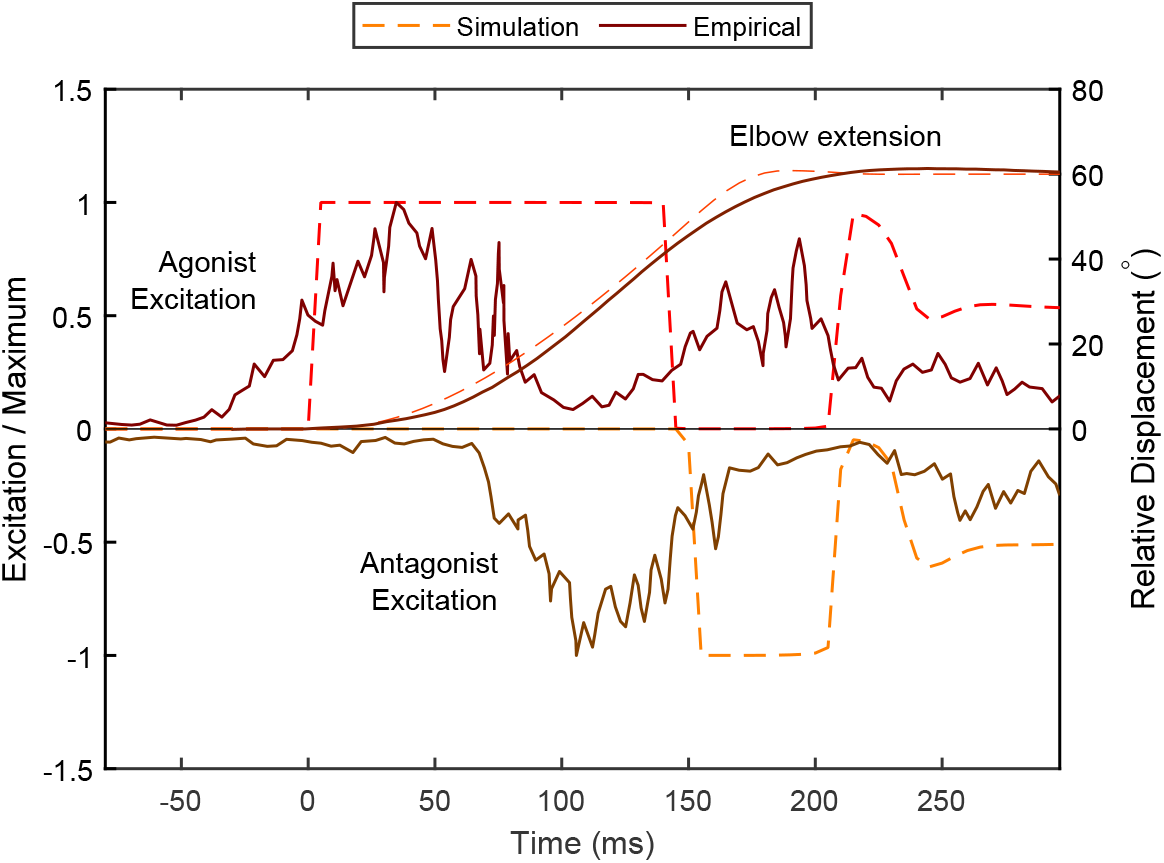
A predictive simulation with baseline parameters exhibits qualitative similarities to empirical EMG data from Forgaard et al. (2013). Both exhibit a smooth arm angle profile with similar maximum velocity, and a triphasic pattern of muscle excitation: a peak in the agonist excitation is followed by a peak in the antagonist and a subsequent peak in the agonist. In addition, both model and empirical data have a sustained level of co-excitation after the arm has come to a stop. EMG peaks occur earlier than those predicted by the model, and have a relatively smooth rise and fall. Empirical data were extracted using WebPlotDigitizer (Rohatgi, 2024).

When input parameters change from baseline, differences in optimal trajectories emerge, although the general pattern remains the same. In the baseline case, muscle length overshoots the target before rapidly settling to the target value (Figure 3A, red line). With the “youngest” parameter combination (highest *F*_max_ and *v*_max_, and lowest *α*_*a*_, *α*_*d*_ and *c*_1_) as well as the “oldest” (lowest *F*_max_ and *v*_max_, and highest *α*_*a*_, *α*_*d*_ and *c*_1_) the same pattern emerges, but with about 2/3 and twice the time to reach the target, respectively, and no overshooting in the “oldest” condition. The velocity profiles resemble a skewed “bell-shape” (Wong et al., 2021), with peak velocity occurring late in the propulsive phase (Figure 3B). Velocity fluctuations during stabilisation increase from the “older” case, to baseline, to the “younger” case. Agonist activation increases early in the movement, and begins deactivating as agonist activation begins (Figure 3C), reflecting the triphasic pattern. The “youngest” condition achieves full activation, and exhibits higher compensatory antagonist activation. In the baseline case, and even more so in the “oldest” case, peak activation is smaller. Agonist activation leads to an initial rapid increase in agonist force (Figure 3D), which decays due to the increase in contractile velocity. When muscle length is close to the target, the antagonist produces a large braking force, well above the agonist peak force, which is then followed by an increase in agonist force as the system slows down. Both agonist and antagonist settle into an equal and opposite sustained force until the end of the simulation time (0.4 s).

**Figure 3:**
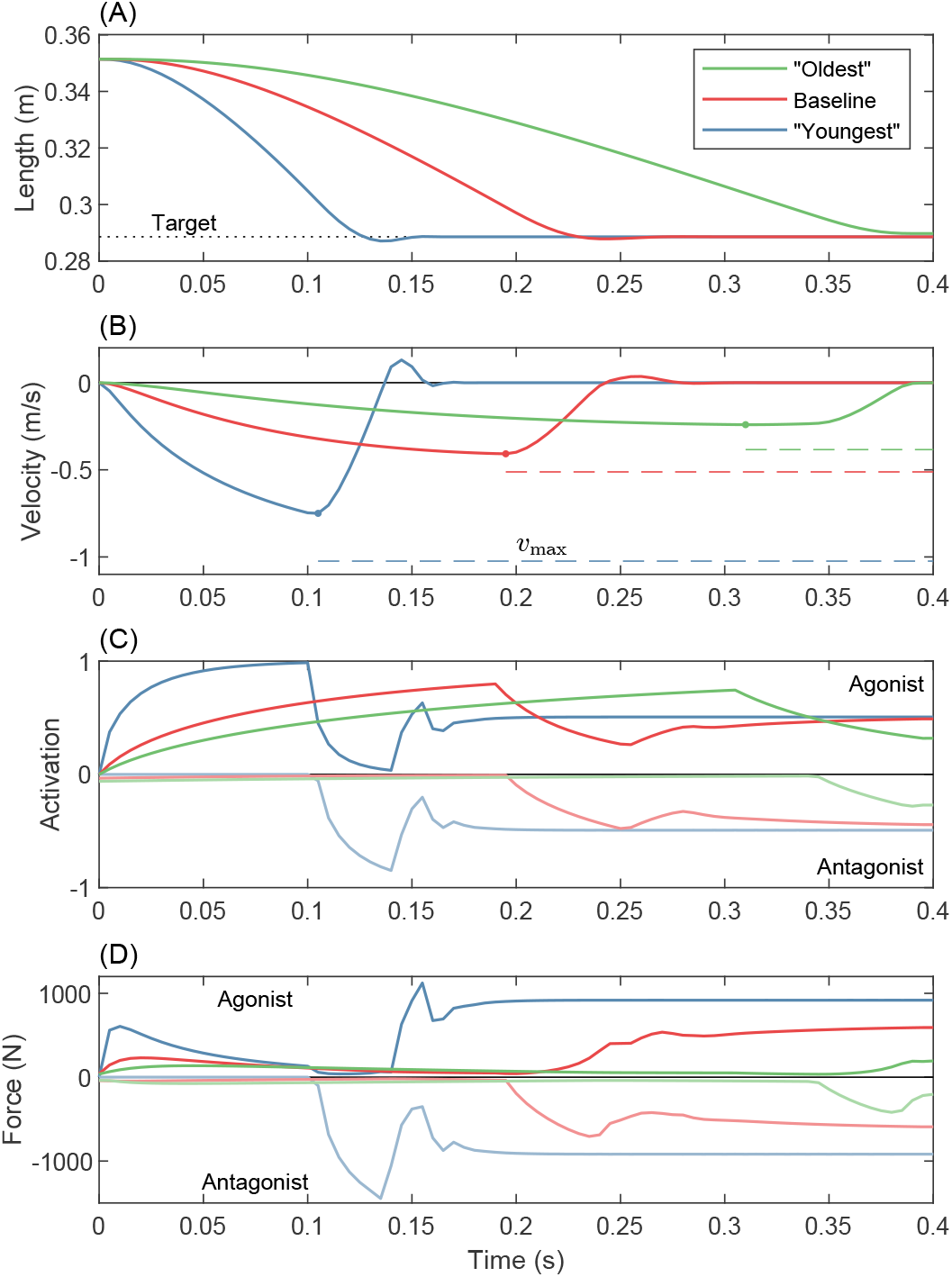
Time series results of optimal solutions with varying input parameters. The “oldest” condition (green) represents muscle with maximal stiffness (*c*_1_ = 0.09), maximum activation and deactivation time constants (*α*_*a*_ = 189 ms, *α*_*d*_ = 130 ms), minimal peak force (*F*_max_ = 650 N), and minimal peak strain (*v*_max_ = 1.2 *l*_0_ s^-1^); i.e. the bottom right corner of the bottom right panel in Figure 4. The “youngest” condition (blue) represents the opposite extreme; the top left corner of the top left panel in Figure 4, with *c*_1_ = 0, *α*_*a*_ = 14 ms, *α*_*d*_ = 10 ms, *F*_max_ = 1950 N and *v*_max_ = 3.2 *l*_0_^-1^. The baseline case (red) represents intermediate values, outlined in Table 2. Note that “youngest” and “oldest” represent extremes explored in the parameter sweep, and not necessarily the expected change in muscle properties through a person’s lifetime. (A) Agonist muscle length reaches the target in all cases, but more quickly for the “younger” muscles. (B) Agonist contraction velocity increases more rapidly in the “younger” muscle. The peak contraction velocity (dot marker) is closest to *v*_max_ (dashed line) in the baseline case. (C) The “youngest” agonist rapidly approaches peak activation, whereas the “older” muscles reach lower levels of peak activation before deactivating for the braking phase. Antagonist muscle (“negative” activation) shows similar patterns. (D) “Younger” muscle displays higher levels of acceleratory and braking force, as well as higher levels of sustained cocontraction after stabilisation.

A performance landscape across the range of parameter combinations is shown in Figure 4. In general, parameters move from “younger” to “older” values from the top left to bottom right– in other words, some combination of increase in parallel stiffness (upper four panels to lower four panels), decrease in *v*_max_ (top row to bottom row for a given stiffness), decrease in force (left to right column of panels), or increase in deactivation or activation time (downward or rightward in a single panel, respectively). As parameters shift to “older” values, performance generally decreases– corresponding to an increase in root-mean-square-error (RMSE). However, there are two notable exceptions. In the absence of stiffness (Figure 4A; panels i-iv), changing deactivation time appears to have no effect on performance (indicated by vertical contour lines). When stiffness is appreciable (Figure 4B; panels v-viii), increasing deactivation time decreases performance. However, for any point in A, the corresponding point in B may have relatively higher or lower RMSE, depending on its location.

**Figure 4:**
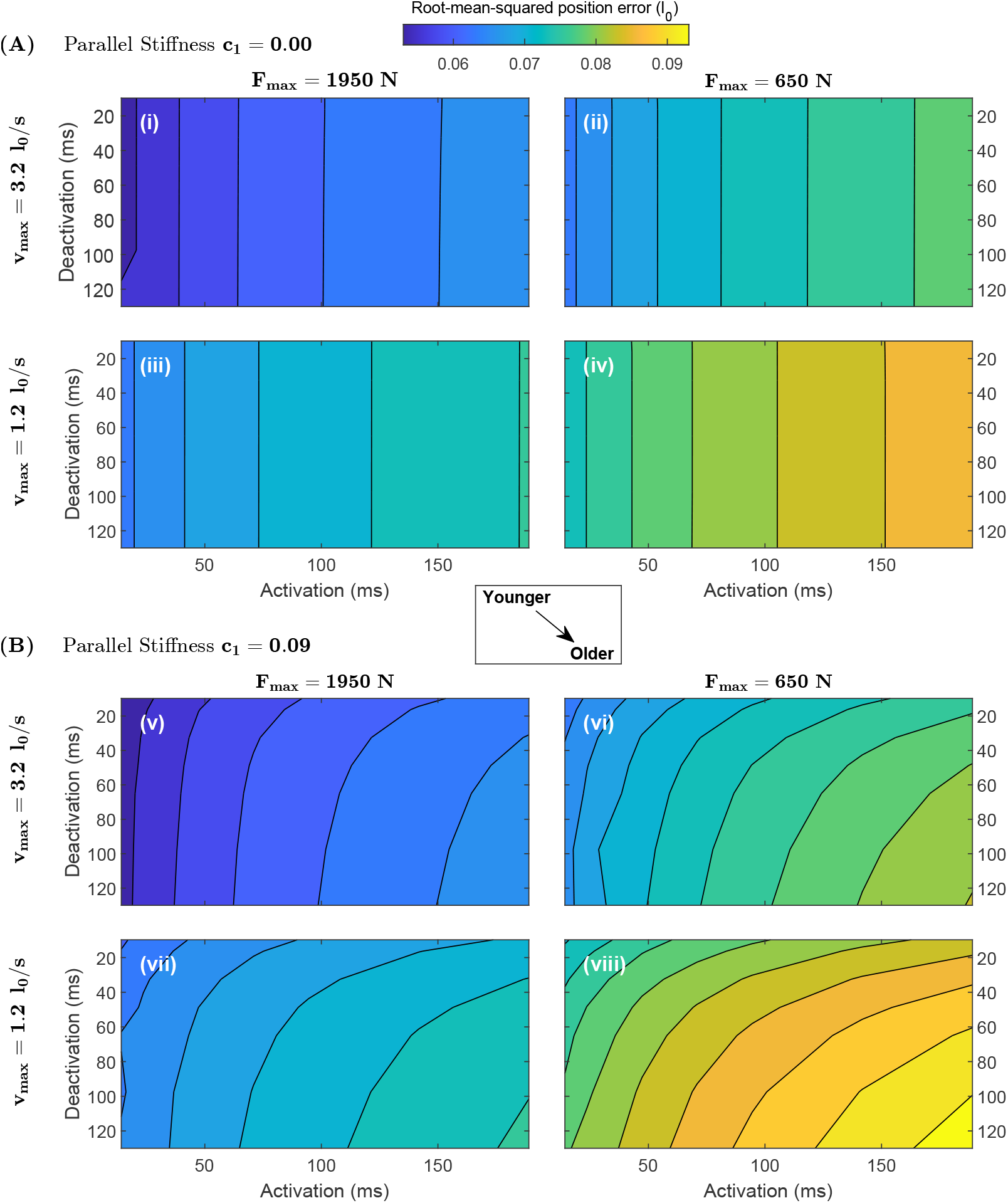
Contours of total Root-Mean-Squared Error (RMSE) during ballistic arm pointing tasks show generally decreased performance as parameters shift towards “older” values, and an interaction between deactivation rate and stiffness. Each panel shows RMSE as a function of deactivation and activation time (increasing to the bottom right), for some combination of maximum strain rate (*v*_max_), peak active tension (*F*_max_) and passive parallel stiffness (*c*_1_). The top four panels (i-iv) are for no passive parallel stiffness, and the bottom four (v-viii) for high stiffness (*c*_1_ = 0.09). Within each stiffness condition, *F*_max_ decreases from the left column to right column of panels, and *v*_max_ from the upper row to the lower row. Altogether, the plots are arranged such that moving from top left to bottom right indicates a shift from relatively “young” to relatively “old” muscle (A) If the muscle has no passive stiffness (*c*_1_ = 0), RMSE increases (*i.e.* performance is reduced) when activation time increases, *F*_max_ decreases, and/or *v*_max_ decreases, but the vertical contour lines indicate that performance does not vary with changes to deactivation time. (B) When the muscle has high stiffness (*c*_1_ = 0.09), RMSE also increases with deactivation time. However, relative to the same parameter combinations without muscle stiffness (the equivalent point in A), the RMSE can either decrease (rightward shift in contour lines at low deactivation) or increase (leftward shift in contour lines). Parameter ranges are larger than the expected natural variation with age, especially in activation and deactivation; see Methods.

This phenomenon is illustrated in Figure 5, showing a performance landscape as a function of deactivation time and stiffness; all other parameters are held constant at baseline values. A hypothetical boundary across the parameter space emerges (Figure 5, white dash line). When deactivation time values are near this boundary, stiffness has little to no effect on performance. When deactivation time is higher than these values (below the line), increasing stiffness decreases performance (indicated by an increase in RMSE). When deactivation time is lower (above the line), increasing stiffness *increases* performance.

**Figure 5:**
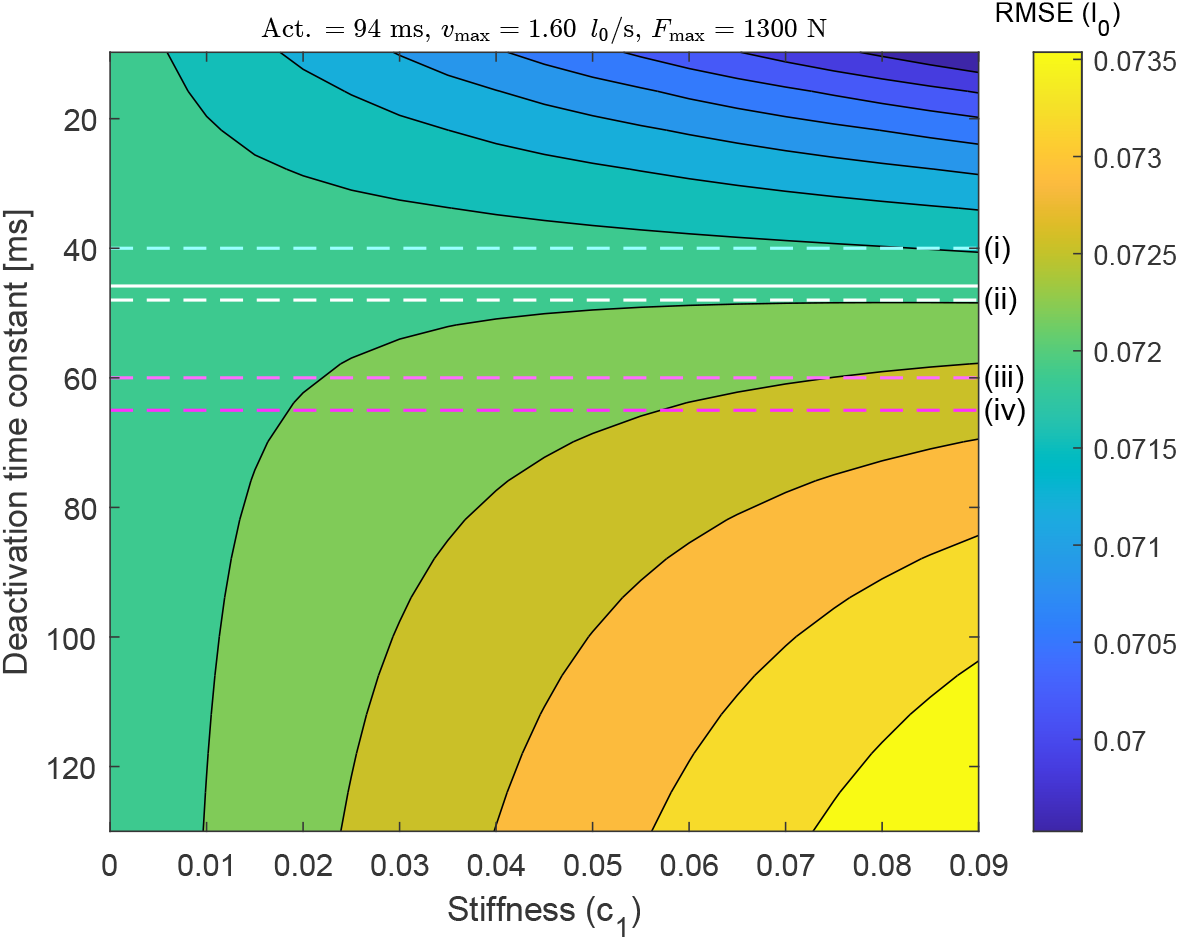
Root Mean Squared Error (RMSE) contours as a function of deactivation time and passive parallel stiffness, with other parameters held at baseline values. When the deactivation is fast (upper portion of plot; small time constant), increasing stiffness decreases RMSE and enhances performance. When the deactivation is slow (large time constant), increasing stiffness decreases performance. Intermediate values of deactivation exist where stiffness changes have little to no effect; at deactivation time of 46 ms (white line), a 0.01 unit change in *c*_1_ changes RMSE by less than 10^−2^ *l*_0_. Various studies use this same activation model in dynamic simulations of ageing, but with different deactivation time constants– potentially leading to different effects of passive stiffness (dashed lines). (i) The default OpenSim value (e.g. used by Karimi et al., 2021) is 40 ms, implying an increase in performance with increased stiffness. (ii) Nowakowski et al. (2022) used an aged value of 48 ms in their simulation, which would lead to no stiffness effect in the current study. (iii) Thelen (2003) used 60 ms and (iv) Murtola and Richards (2022) used 65 ms in their aged models. These values would lead to a decrease in point-to-point performance with increased stiffness, as found by Murtola and Richards (2023) with a related model.

In the presence of stiffness, muscle deactivation affects the net accelerative torque about the joint. In the initial posture, the agonist is stretched, increasing its passive contribution (Figure 6A, *t* = 0). To maintain stasis prior to joint movement onset, the antagonist must supply an equivalent active force (Figure 6B). As the agonist contracts and the elbow extends, the antagonist becomes stretched, experiencing force amplification as it shifts to the eccentric part of the force velocity curve. If the antagonist can “switch off” quickly, it can reduce its resistive force below the agonist’s passive contribution. In this case, the net extending torque about the joint is larger than what it would be without any passive stiffness (Figure 6C, cool colours), and the passive stiffness has a net benefit to performance. However, if the antagonist cannot switch off quickly, then its force rises *above* the agonist’s passive contribution (Figure 6B, warm colors), and the net extensor torque is reduced compared to the no-stiffness case (Figure 6C, warm colours).

**Figure 6:**
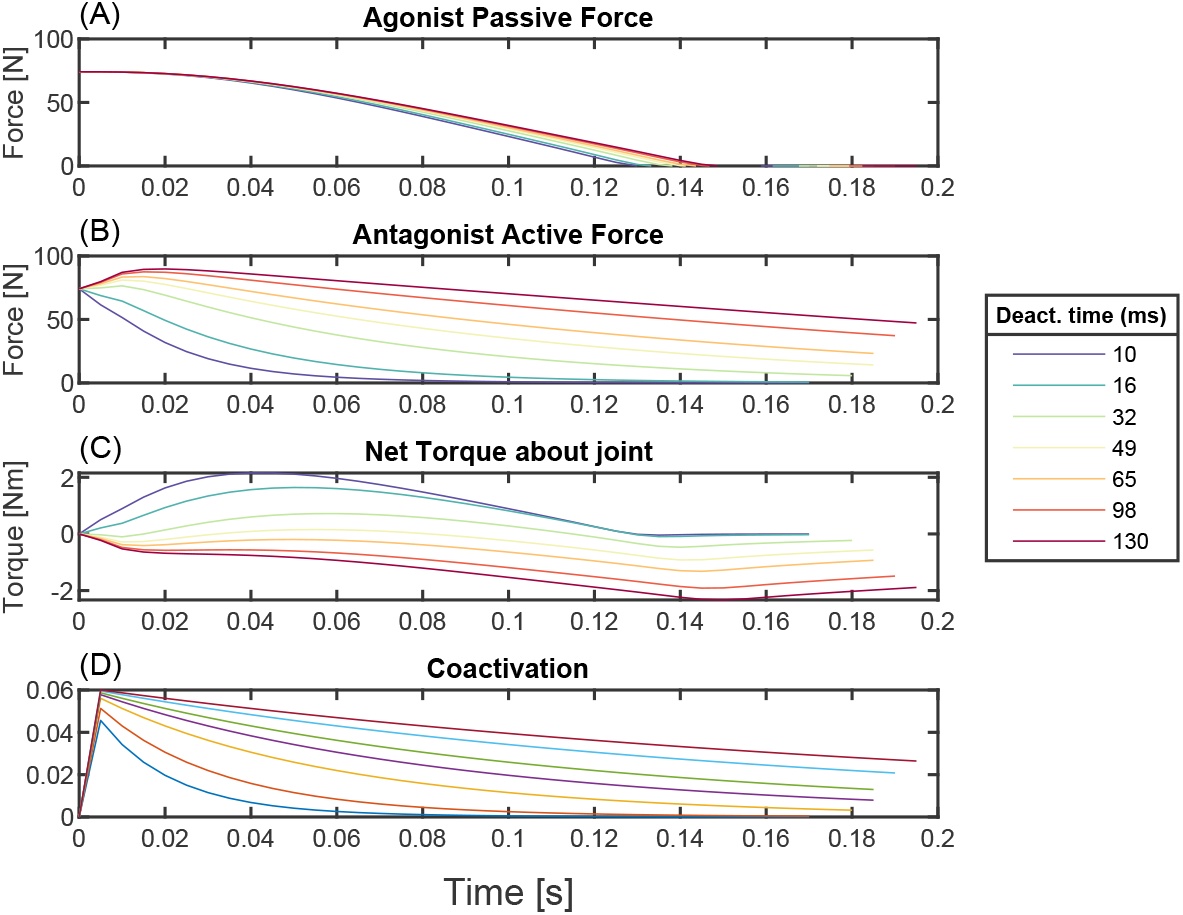
Deactivation time affects the net force about the joint in the presence of agonist passive parallel force. Shown here are results with baseline parameters in all but deactivation time, and *c*_1_ = 0.09 (A) Agonist passive force is positive at *t* = 0 as the muscle is stretched, generating a positive torque about the joint. (B) The antagonist must produce an equal and opposite active force to keep the joint at rest at *t* = 0. As the arm begins to accelerate, the antagonist is stretched, moving it to the eccentric part of the force velocity curve (Figure 1C). If the muscle can deactivate quickly (cooler lines), its force can rapidly drop. However, if the muscle deactivation is slow (warmer lines), the antagonist’s force rapidly increases in excess of the extra passive force from the agonist. (C) The net torque at the joint, in addition to the active force from the agonist, thus depends on activation level. If deactivation time is low, the system gets an accelerative boost from the antagonist’s passive force. If the deactivation time is high, the antagonist’s resistive torque exceeds the passive torque, and the additional net torque is negative. (D) Early-onset coactivation occurs regardless of deactivation time. However, coactivation persists at higher levels when deactivation time is higher (warmer lines)

Because the antagonist is initially active, coactivation is high at the beginning of the acceleration phase and decreases in time, more quickly for fast deactivation (Figure 6D). Coactivation also increases with deactivation in the absence of stiffness, but only in the the braking phase. Figure 7 shows the results of changing deactivation while keeping stiffness zero and other parameters at baseline. The agonist muscle strain trajectories are virtually indistinguishable (Figure 7A), with only slight differences in oscillation around the target point while the system comes to a stop. The strain achieves similar peak velocities (Figure 7B), with lower deactivation times resulting in higher amplitude, higher frequency oscillations about zero during the braking phase. The onset of agonist deactivation occurs slightly earlier when deactivation time is larger (15 ms earlier for the highest *versus* lowest *α*_*d*_), and the antagonist begins activating slightly earlier as well (at most 5 ms). As the arm slows down, antagonist deactivation remains high when deactivation is slower, and the agonist increases its activation for longer. This results in larger sustained forces in both muscles (Figure 7D), leading to higher levels of cocontraction both during the braking phase and once the simulation has terminated (Figure 7E). All deactivation conditions have sustained levels of equal activation and force in both muscles at the end of the motion.

**Figure 7:**
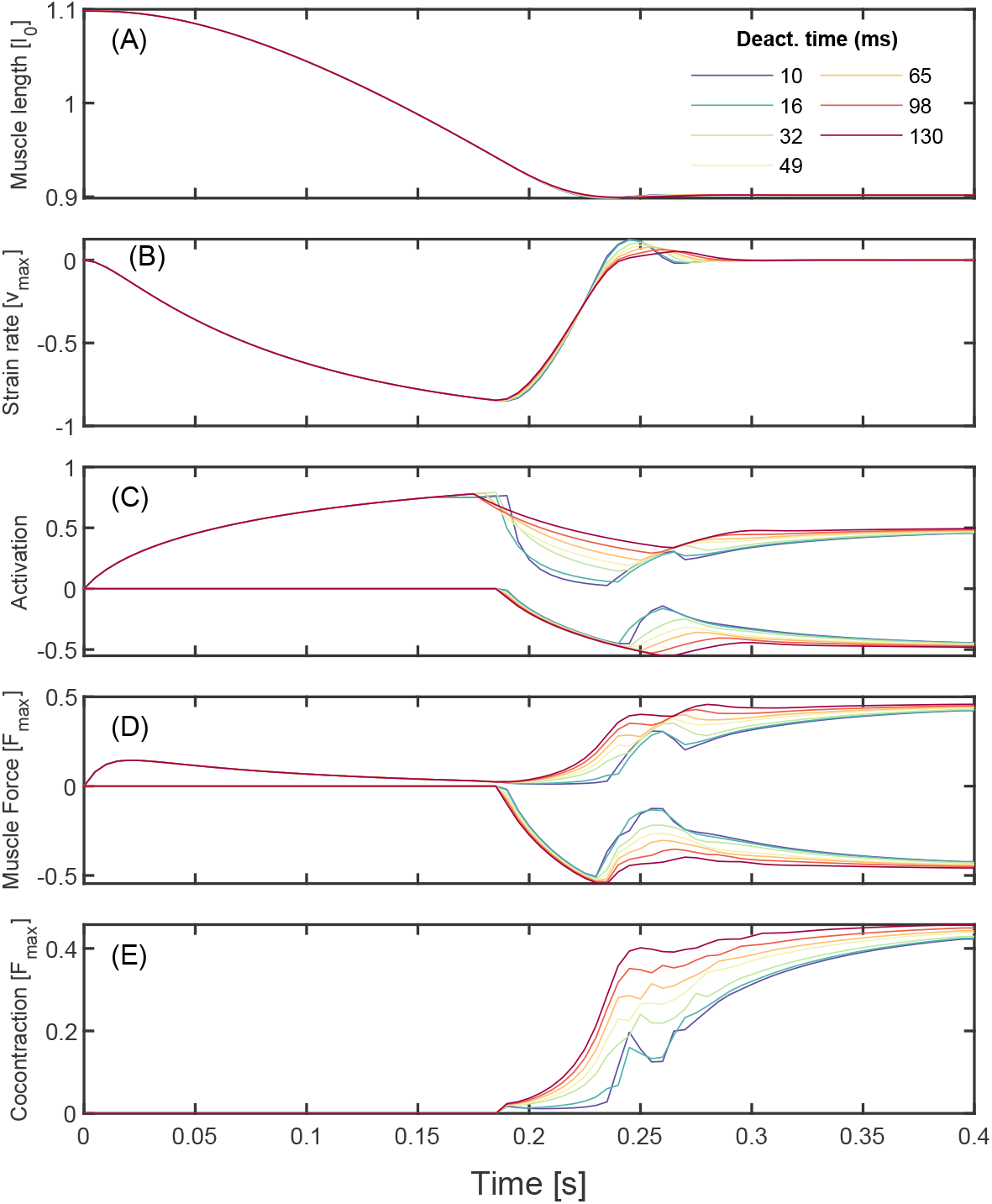
Slowing deactivation increases the level of cocontraction in optimal ballistic movements at zero parallel stiffness, without affecting performance. Here all other parameters are held constant at baseline values (Table 1). (A) Agonist muscle length trajectories appear nearly identical for all deactivation times. (B) Slight differences in agonist muscle velocity emerge during the deceleration and stabilisation portion of the trajectory. Muscle contraction velocity approaches *v*_max_ in all cases. The antagonist strain and strain rate are equal and opposite at every instant. (C) Positive activation corresponds to the agonist, while negative corresponds to the antagonist. When deactivation time is low (cool colours), the agonist rapidly decreases activation as the antagonist activation rises. When deactivation time is high (warm colours), the agonist cannot deactivate fast enough, and the antagonist exhibits a corresponding higher level of peak activation, resulting in higher sustained levels of activation post stabilisation. (D) Because the agonist’s muscle velocity approaches *v*_max_, the agonist’s force is small immediately before the target is reached. (E) Cocontraction initiates in the braking phase (later than in the conditions with passive parallel stiffness; Figure 6), and achieves higher sustained levels post stabilisation with slower deactivation.

In order to determine the biggest contributors to performance in this dataset, we performed linear regression of RMSE against the five variables and their interactions plus a constant (a total of 16 terms). Best-fit coefficients are shown in Table 3, where the magnitude of the coefficient represents how much changing the specific term from its minimum to maximum value (holding all other terms and interactions constant) changes RMSE on average (as a fraction of total possible change). By analogy to the “standardised beta” statistical technique (Mizumoto, 2023), we define these coefficients as the “importance” of the term. Five of the 16 coefficients are important to first order (magnitude *>* 10^−1^). One is the intercept value, and three others are the individual parameters of activation rate, maximum strain rate and maximum isometric force; deactivation rate and stiffness are not important on their own to first order. The only important interaction effect (to first order) is that of deactivation and stiffness together.

**Table 3:** Results of linear model fitting, including individual and interaction terms. Each term in the model is normalized to its range (Table 2). Coefficient estimates represent the relative “importance” of that term. Estimates and standard error are shown to one significant digit. Bolded values represent a coefficient estimate with magnitude *>* 10^−1^: *i.e.* effects that are “important to first order”. Note that all individual coefficients are important to first order except for activation and stiffness; however, together these have the only interaction coefficient with magnitude *>* 10^−1^.

| Corresponding<br>Term | Coefficient estimate |  |  |
| --- | --- | --- | --- |
|  | All<br>Terms | First-order<br>terms only | First-order<br>independent<br>terms only |
| Intercept | **5e-1** | 5e-1 | 5e-1 |
| $\alpha_d$ | 2e-2 | | |
| $c_1$ | 6e-3 | | |
| $\alpha_a$ | **3e-1** | 3e-1 | 3e-1 |
| $v_{\max}$ | **-2e-1** | -2e-1 | -2e-1 |
| $F_{\max}$ | **-2e-1** | -3e-1 | -3e-1 |
| $\alpha_d : c_1$ | **1e-1** | 8e-2 | |
| $\alpha_d : \alpha_a$ | 3e-2 | | |
| $\alpha_d : v_{\max}$ | -3e-2 | | |
| $\alpha_d : F_{\max}$ | -4e-2 | | |
| $c_1 : \alpha_a$ | -2e-2 | | |
| $c_1 : v_{\max}$ | -6e-3 | | |
| $c_1 : F_{\max}$ | -6e-2 | | |
| $\alpha_a : v_{\max}$ | 4e-2 | | |
| $\alpha_a : F_{\max}$ | -6e-2 | | |
| $v_{\max} : F_{\max}$ | 5e-5 | | |
| $R^2$ | 0.95 | 0.94 | 0.92 |

To further check whether these first-order terms account for the vast majority of the variation, we fit a linear model with only these terms (Table 3). The fit has R^2^=0.94, 1% smaller than if all 16 terms are included, indicating that first-order terms are sufficient to explain most of the variation in performance. A fit further removing the interaction term yields R^2^=0.92.

We also performed linear regression of *J*_*RMSE*_ against mean coactivation in each simulation, to see whether coactivation was associated in any way with performance. Mean coactivation (*ā*_*c*_. equation 4) ranges from 0.034 to 0.415. We find that mean coactivation is strongly correlated with performance (R^2^ = 0.7), and that higher coactivation is associated with better performance (*J*_*RMSE*_/ max(*J*_*RMSE*_) = −0.89 *ā*_*c*_ + 0.93, *p <* 10^−6^).

## 5 Discussion

The aim of this study is to determine how age-related changes to muscle contractile properties affect ballistic reaching performance in a point-to-point task. Here performance is defined as the cumulative root-mean-square distance from the arm to the target (Equation 3), and is driven by how quickly the arm can both reach and converge to its target point. When control is optimised to maximise performance, it resembles rapid point-to-point movement in human elbow flexion (Forgaard et al., 2013, Figure 2), including displacement trajectories that are similar to empirical data in shape and duration, and a triphasic excitation pattern– a well-documented phenomenon in ballistic movements (Leib et al., 2020; Correia et al., 2022). This gives us confidence that the simplified model captures salient features of rapid targeted joint excursion. By systematically changing parameters and re-solving the optimisation problem, we constructed a performance landscape as a function of muscle parameters to determine how shifts in muscle parameters affect optimal higher performance (Figure 4).

We find that reductions in peak isometric force and maximum strain rate– all associated with ageing muscle– each lead to decreases in performance, and that these effects do not interact. Increases in both deactivation time and stiffness– also associated with ageing– tends to decrease performance as well (although the exact effect depends on their interaction). To our knowledge, this is the first study to show that each of these five age-related changes to muscle can deleteriously affect performance.

Because of the model’s relative simplicity, we can interrogate the solutions to understand the mechanistic links between parameter change and performance. In the following, we propose mechanistic reasons behind key findings: why activation time, peak force and maximum strain rate are the main contributors to performance; why deactivation time and stiffness interact; and why cocontraction, perhaps surprisingly, is associated with higher speed and overall performance in this model.

### 5.1 Activation time, maximum force, and maximum muscle velocity are the main contributors to performance

Figure 4 shows that increases in activation time, decreases in peak force or decreases in maximum strain rate lead to a reduction of performance (when holding other parameters constant). A linear fit of these three variables alone (without interactions but including the intercept; Table 3) explains 92% of the variation in performance, encapsulating their dominance in determining performance. The intercept value remains in place because the displacement error will remain at a constant value if the arm cannot move. The remaining 12 terms explain only an additional 3% of the variation in performance in the linear model. Our results suggest that age-related reductions in activation rate, maximum muscle strain rate, and isometric force all contribute overwhelmingly to reductions in ballistic performance. Because of the independence of these effects, we infer that a detectable reduction in any of these parameters with age generally has a negative effect on rapid point-to-point movement.

As the performance metric of RMSE is driven largely by time to target (Figures S1 and S2), these results come readily by considering each term’s contribution to the time-averaged applied force. Since the load and initial distance to target are constant, maximising time-averaged propulsive force is equivalent to minimising time. A lower activation time constant increases average force by allowing the muscle to reach full activation more quickly. A higher *F*_max_ increases the force available from full activation; and a higher *v*_max_ means the applied force is attenuated less as velocity increases.

In contrast, deactivation time and stiffness do not, on their own, affect performance in this dataset. They do, however, interact to affect performance, and only with each other (to first order; Table 3). These results may seem at first counterintuitive. Since the arm must slow down to reach the target, it may seem like slow deactivation would be detrimental– analogous to trying to stop a car with the accelerator pressed down. Moreover, since passive stiffness increases the applied force of the antagonist during acceleration and the agonist during braking, it may seem like increased passive stiffness should always be beneficial. Both these counterintuitive results can be explained by the force-velocity properties of muscle, as we discuss in the following sections.

### 5.2 In the absence of passive stiffness, deactivation rate does not affect ballistic performance

If the agonist cannot turn off quickly during the braking phase, then it supplies excess force as the joint approaches its target. In order to stop in time, the controller must use one or a combination of three strategies: (1) the agonist reduces approach speed by reducing its applied force, (2) the antagonist applies braking force sooner, and/or (3) the antagonist applies extra braking force. The first strategy occurs in our simulations, but the effect is minor. As deactivation time increases, the agonist begins switching off only slightly earlier (by at most 15 ms; Figure 7B). The second strategy also occurs, but is likewise minor; as deactivation time increases, the antagonist comes online slightly earlier, but by at most 5 ms (Figure 7C). Despite these changes to force timing, the peak approach speed remains virtually unchanged (Figure 7 b), and these small changes do not account for the similarity in performance.

The third strategy is much more effective– and is enabled by the muscle’s force-velocity properties. Because both muscles have identical properties, and the antagonist operates on the eccentric part of the force-velocity curve (Figure 1A,C), it can always produce much larger force than the agonist as the joint is slowing down. Therefore, if the agonist remains stuck “on” due to slow deactivation, the antagonist can simply maintain heightened activation to overpower the agonist force (Figure 7D), leading to higher cocontraction (Figure 7E). The higher cocontraction effectively increases active damping at the joint (Seidler-Dobrin et al., 1998; Latash, 2018): if the antagonistic pair coactivates, a higher velocity is met with a higher force resisting motion, and this resistive force increases with the level of coactivation. This coactivation-dependent active damping compensates for the sustained propulsive force at large deactivation times. Therefore, slower deactivation rates lead to higher levels of cocontraction but not to lower performance– at least in the absence of stiffness.

### 5.3 Stiffness and deactivation interact to enhance or reduce performance due to antagonist release

Increasing agonist stiffness can enhance performance, but only if the antagonist deactivation time is sufficiently small (Figure 5). This occurs because the initial flexed posture of the joint requires the agonist to be stretched, and therefore to supply passive torque about the joint (Figure 6A). To maintain stasis, the passive torque must be matched by active force from the antagonist (Figure 6B). Movement initiation from the agonist stretches the antagonist, shifting it to the eccentric part of the force-velocity curve. If the deactivation time is low enough, the antagonist force can drop below the agonist passive force, and the net contribution to joint torque is positive (Figure 6C, cooler lines). If instead the deactivation time is high, the antagonist force quickly exceeds the passive contribution– there is then a net decrease in torque about the joint (Figure 6D warmer lines). Therefore, increasing stiffness can have a net positive or negative effect on performance, depending on the deactivation rate (Figure 5). If deactivation is slow, the antagonist slows the joint down, whereas if deactivation is high, the increased agonist stiffness provides extra positive torque about the joint, in excess of the antagonist’s active force.

More specifically, our results suggest a critical deactivation time (approximately 45 ms) where increased stiffness has no effect on performance. In our model, if deactivation time is greater than this value, then age-related increases in stiffness have a detrimental effect on performance; whereas if deactivation time is smaller, then increased stiffness enhances performance. This sensitivity to deactivation time has implications for dynamic simulation studies of musculoskeletal ageing. For example, the default OpenSim (Seth et al., 2018) deactivation time constant is 40 ms, used by Karimi et al. (2021) in a model of walking in the elderly. Using this value within our model would result in a slight positive effect of increased stiffness. Nowakowski et al. (2022) increased *α*_*d*_ to 48 ms in their aged condition to assess fall risk in an OpenSim model. This value would indicate little to no effect of increased stiffness in our model. Thelen (2003) and Murtola and Richards (2022) used deactivation times of 60 and 65 ms respectively in their simulations of musculoskeletal ageing, with the latter finding a decrease in performance with increased stiffness, in accordance with Figure 5. Because of the sensitive interaction between passive stiffness and deactivation time, care should be taken in the selection of both parameters when inferring age-dependent effects.

Both increased deactivation rate and increased stiffness are associated with higher levels of coactivation and cocontraction. However, in our dataset cocontraction is not a feature to be avoided, but rather a behavioural outcome of maximising rapid movement performance.

### 5.4 Cocontraction can be an optimal strategy for rapid movement

Arm reaching in elderly individuals is associated with higher levels of cocontraction (Darling et al., 1989; Seidler-Dobrin et al., 1998; Wittenberg et al., 2022), and is generally slower, both in reaction time and total movement time (Bautmans et al., 2011; McKinnon et al., 2017; Summerside et al., 2024). Ageing is also associated with reductions in neural conduction velocity (Xi et al., 1999; Di Iorio et al., 2006), which may lead to feedback delays. Since cocontraction increases joint stability against perturbations without requiring feedback control (Gribble et al., 2003; Latash, 2018), it has been suggested that elderly persons cocontract to stabilize their joints against perturbations (Seidler-Dobrin et al., 1998), but that coactivation comes at a price of slowing movements (Bautmans et al., 2011; Lo et al., 2017; Arpinar-Avsar and Celik, 2020; Wittenberg et al., 2022) and reducing power production in older adults (McKinnon et al., 2017).

The present results offers another view. The current model has no outside perturbation, and no control deficits-the controller has an exact representation of the current states and system dynamics, and can change excitation to any value at 5 ms temporal resolution. And yet, cocontraction emerges naturally as an optimal strategy to maximize performance– which in this case is driven largely by minimizing time to target (Figure S2). Indeed, trials with higher levels of coactivation tended to reach the target and stabilise *more quickly* than trials with lower levels of coactivation (Figure S3).

In the idealised system under investigation, the controller provides an optimal sequence of muscle excitations that result in certain amounts of coactivation. By definition, any increase or decrease in coactivation level from this optimum would decrease performance. Our results suggest that, for a given musculoskeletal system, there exist optimal levels of coactivation and cocontraction for rapid point-to-point movements, which depend on the physiological and physical properties of the system. This implies that differences in coactivation between two biological systems (e.g. young and old participants) do not indicate *a priori* whether either system is operating optimally for that task– the optimal level of coactivation depends on physics, physiology and the objective of the movement.

Our data suggest several heuristics for how and when coactivation is associated with performance differences. During the propulsive phase, the association between cocontraction and performance depends on the interaction between agonist stiffness and antagonist deactivation rate. Cocontraction will occur when the antagonist initially resists the agonist passive force. For a given level of stiffness, increased cocontraction is correlated with a loss of performance, as it indicates that the antagonist cannot deactivate quickly enough (Figure 6); but for two systems with different stiffnesses (with all other parameters similar), increased cocontraction may be associated with an increase or decrease in performance, depending on the muscle deactivation rate.

During the braking phase, increased deactivation time leads to increased cocontraction (Figure 7E). If all other parameters are equal, however, this does not affect performance, because cocontraction increases joint damping that compensates for relatively increased agonist activity. After stabilisation, optimal solutions with higher agonist activation during propulsion generally remain at higher levels of coactivation (Figure 3). In this dataset, because higher-performance trials tended to have shorter ballistic phases and longer periods after braking, higher-performance tends to be associated with higher levels of coactivation (Figure S3). Empirical evidence aligns with this finding; Suzuki et al. (2001) found that faster point-to-point arm movements exhibited higher levels of post-movement coactivation.

In human and animal ballistic data, it is important to carefully consider how the absence of cocontraction would affect the performance dynamically, before claiming a causal link between the two. We have shown that, even in a simple, two-muscle model, the effects of cocontraction on movement time can be subtle and not immediately intuitive. The emergence of coactivation and cocontraction in this model, without any feedback delays or perturbations, suggest that increased coactivation in ageing may arise in part from increases in muscle stiffness and deactivation time, and that coactivation is not, in of itself, responsible for reductions in movement speed in the elderly.

This does not, however, preclude cocontraction as a compensatory strategy to reject perturbations without feedback; both a loss of rapid feedback control and the above physiological changes to muscle might lead to increased cocontraction in the elderly. The underlying causes of increased coactivation with age, and its associations with movement performance, will be clarified with further experiments, as well simulations that incorporate perturbation, signal noise or signal delays into the model.

### 5.5 Limitations and future directions

The use of a simplified computational framework allowed us to explore a large space of muscle property changes that are associated with age. We were uniquely able to determine mechanistic links between parameters and performance because all aspects of the simulation are tightly controlled. Such analysis would be much more difficult, if not impossible, *in vivo* because ageing is a multifactorial condition wherein not all relevant parameters can be accurately measured, inferred or known.

Our study used a Hill-type model because such models are well-studied, widely used in reaching simulations (Louis and Gorce, 2009; Yu and Wilson, 2014; Murtola and Richards, 2022) and present low computational cost, making them well-suited for large parameter sweeps such as in the current study. However, more sophisticated muscle models (Nishikawa et al., 2011; Schappacher-Tilp et al., 2015) could be used in future work to address transient and hysteresis effects. Future models could also change task parameters by introducing perturbations or including energetic cost in the objective (Wong et al., 2021; Summerside et al., 2024). Further complicating features such as joint damping, variable moment arms, or differences between muscles could be implemented in follow-on studies, at the expense of computational cost.

The influence of parameters on performance was explored using linear regression based on the standardized beta approach (Mizumoto, 2023). Because predictors are first normalised to their range in the linear regression, the magnitude of their regression coefficients (the “importance”) could change if different parameter variation ranges were used in the analysis. For example, if *F*_max_ were kept at relatively narrow values, it would appear to have less of an overall effect than *v*_max_ and *α*_*a*_, if the latter terms maintained larger variation. Interaction effects could also change; if stiffness values were held at a narrow range of large values, then variation of deactivation time on its own would appear to have an effect on performance (Figure 5). The variation in muscle parameters considered in this simulation study is likely larger than natural variation associated with age in humans. For example, Runnels et al. (2005) find that maximal isotonic knee extensor torque in young men is at most twice the value compared to their oldest cohort; whereas we explore a 3-fold change in maximum force. Thus, the absolute changes in performance in this simulation study should not be taken as representative of the changes expected with age– especially when considering the anatomical simplifications used in our model. The qualitative predictions are more likely to be generalisable, and suggest that changes to activation rate, isometric force and maximum strain rate detrimentally influence maximal performance with increasing age.

The current findings on the interaction between muscle parallel stiffness and deactivation rate warrant future experimental investigation. Reported age-related changes to muscle parallel stiffness can differ between studies and exhibit length-dependent effects (Noonan et al., 2020; Xu et al., 2021). Age-related changes to muscle deactivation rate are poorly studied, and the effects can differ between studies– even in the same isolated muscle preparations (e.g. Brooks and Faulkner, 1988; Moran et al., 2006; Kiriaev et al., 2021). Because of this, and the interactivity of stiffness and deactivation rate, future experiments would be required to predict how they affect performance with age *in vivo*. In our dataset, simultaneous increases in stiffness and deactivation time can decrease or increase performance, depending on their correlation and location in the performance space (Figure 5). Despite the above uncertainty, our analysis shows both effects are hypothetically possible, and provides a mechanism why.

## 6 Conclusions

We used a single-joint, two-muscle model to simulate point-to-point arm movements. Optimising excitations to minimise cumulative squared error to target resulted in ballistic motion with realistic triphasic activation. By altering muscle parameters that covary with age, we determined how parameter variation relates to changes in reaching performance. Reductions in force and maximum strain rate, and increases in activation time resulted in decreased performance, and these effects were largely independent of one another. In contrast, increasing parallel stiffness and deactivation time had little effect on performance on their own. For fast deactivation rates, increasing stiffness increases performance, whereas for slow deactivation rates increasing stiffness decreased performance. Both increased stiffness and deactivation times increased coactivation, but increased coactivation was generally associated with better performance in this dataset. This suggests that coactivation may not, in itself, limit movement speed with age, but may reflect increases in characteristic muscle deactivation time and stiffness– factors that can impair performance. Future studies will more precisely characterise age-related changes in muscle deactivation across species and explore how aged muscle affects perturbation response. In doing so, we move closer to uncovering the fundamental links between muscle physiology, motor control and locomotor performance through the lifespan.

## 7 Data Availability

All code and data supporting this article are available at doi: https://doi.org/10.5281/zenodo.15802371

## Supplemental Information

### 11 Smooth approximations of Hill-type dynamics

Muscle force was implemented as a Hill-type model of the form

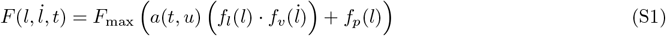

where *f*_*l*_, *f*_*v*_ and *f*_*p*_ are underlying force-length, force-velocity and parallel passive force characteristics, respectively, and *a* is time-dependent activation. These are based on Murtola and Richards (2022, 2023), which in turn follow Otten (1987) and Thelen (2003).

These underlying models introduce non-smoothness which leads to issues in gradient-based optimisation. Therefore, the models were smoothed or adjusted where necessary to avoid these issues, resulting in the smoothed Hill-type model

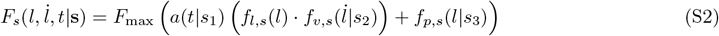

with the smoothing parameter *s*_*i*_ *>* 0. For the results in this paper, we used *s*_1_ = 500 and *s*_2_ = *s*_3_ = 200. The following lays out and defines the smoothed functions used in Equation S2

### S1 Force-length characteristics

Force-length characteristics were specified following (Otten, 1987) as

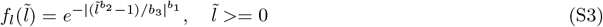

where 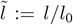 is the normalised muscle length, with *l*_0_ being optimal length. The absolute value function introduces a discontinuity in the derivative; however, if *b*_1_ is even, the absolute value is unnecessary. Therefore, we chose *b*_1_ = 2, instead of 1.3 as in (Murtola and Richards, 2022, 2023), and removed the absolute value function, yielding

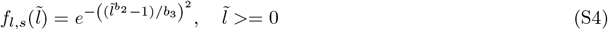

### S2 Force-velocity characteristics

Otten (1987) implements force-velocity as a piecewise function

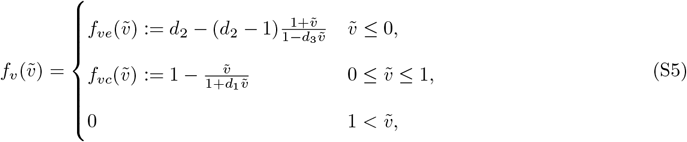

where 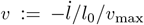 is the normalised contractile velocity. The piecewise nature of the implementation introduces discontinuities, which we removed by sigmoidal smoothing. The sigmoid function

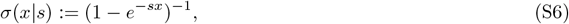

with the smoothing parameter *s >* 0, acts as a smooth approximation of a step function. This allows a piecewise function *f* (*x*) = *{f*_1_(*x*), *x* ≤ 0; *f*_2_(*x*), 0 *< x}* to be approximated as the continuous function *f*_*s*_(*x*|*s*) = *f*_1_(*x*)*σ*(−*x*|*s*) + *f*_2_(*x*|*s*)*σ*(*x*|*s*), with the approximation fidelity increasing with larger values of *s* (at the expense of potentially larger first and second derivatives at the piecewise transition).

Using this technique, we first rewrote Equation S5 as

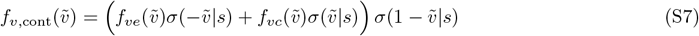

However, as each subfunction *f*_*ve*_ and *f*_*vc*_ is now evaluated everywhere, they introduce singularities when the denominators are zero. These singularities can be mitigated by augmenting the denominator with a smooth ramp function, which ensures the denominator is everywhere positive. The eccentric contraction function 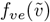 is replaced with the augmented function

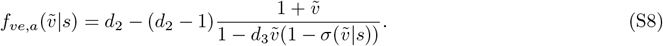

This ensures that when 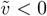 (eccentric contraction), *f*_*ve,a*_≈*f*_*ve*_, and when 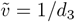 (which would otherwise be a singularity), the denominator evaluates to *σ*(1*/d*_3_|*s*) ≈ 1. For 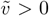, the denominator is always positive, and the eccentric term is attenuated overall by sigmoidal smoothing.

The concentric term is similarly augmented to

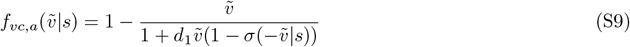

and the fully smoothed approximation of Equation S5 is

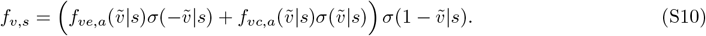

### S3 Parallel Stiffness

Murtola and Richards (2023) developed an exponential model of parallel passive stiffness based on Thelen (2003) and Winters (1995),

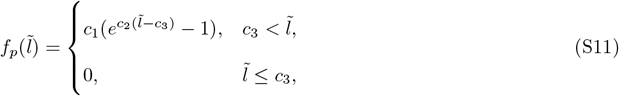

where *c*_3_ is the normalised slack length. Following the procedure outlined above, the piecewise nature of this equation was eliminated through sigmoidal smoothing, generating the smoothed function

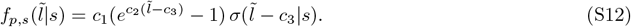

### S4 First-order activation model

The activation model was first-order, based on Thelen (2003). Given the excitation *u*(*t*), the instantaneous activation rate is determined *via* the piecewise differential equation

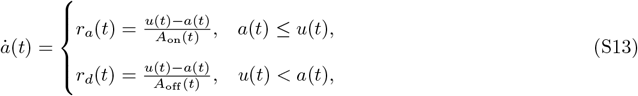

which specifies activation and deactivation rates using time-varying activation and deactivation parameters

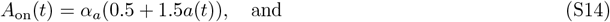

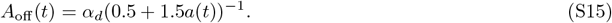

As with other piecewise equations, this was converted to a continuous function with sigmoidal smoothing, yielding the single differential equation

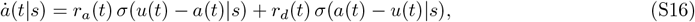

### S5 Correlations between performance metric, time and coactivation

The performance metric of *J*_*RMSE*_ encompasses both a temporal aspect and a spatial aspect. There are two phases to the behaviour: a ballistic phase, achieving rapid movement towards the target, as well as stabilisation phase, where the arm comes to rest at the target.

Here we make the case that the performance metric *J*_*RMSE*_ is driven largely by the time to target. We define time to target *T*_*T*_ as as the first timepoint where the muscle length is within 0.5% *l*_0_ of the target length. We can then calculate the fraction of *J*_*RMSE*_ due to the ballistic phase, compared to the stabilisation phase, as

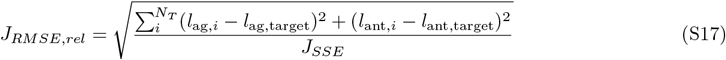

where *N*_*T*_ is the timestep corresponding to *T*_*T*_.

The fraction of *J*_*RMSE*_ accrued during the ballistic phase is never less than 99%. This indicates that the performance metric is largely driven by the speed with which the simulation can reach the target (Figure S1)

**Figure S1:**
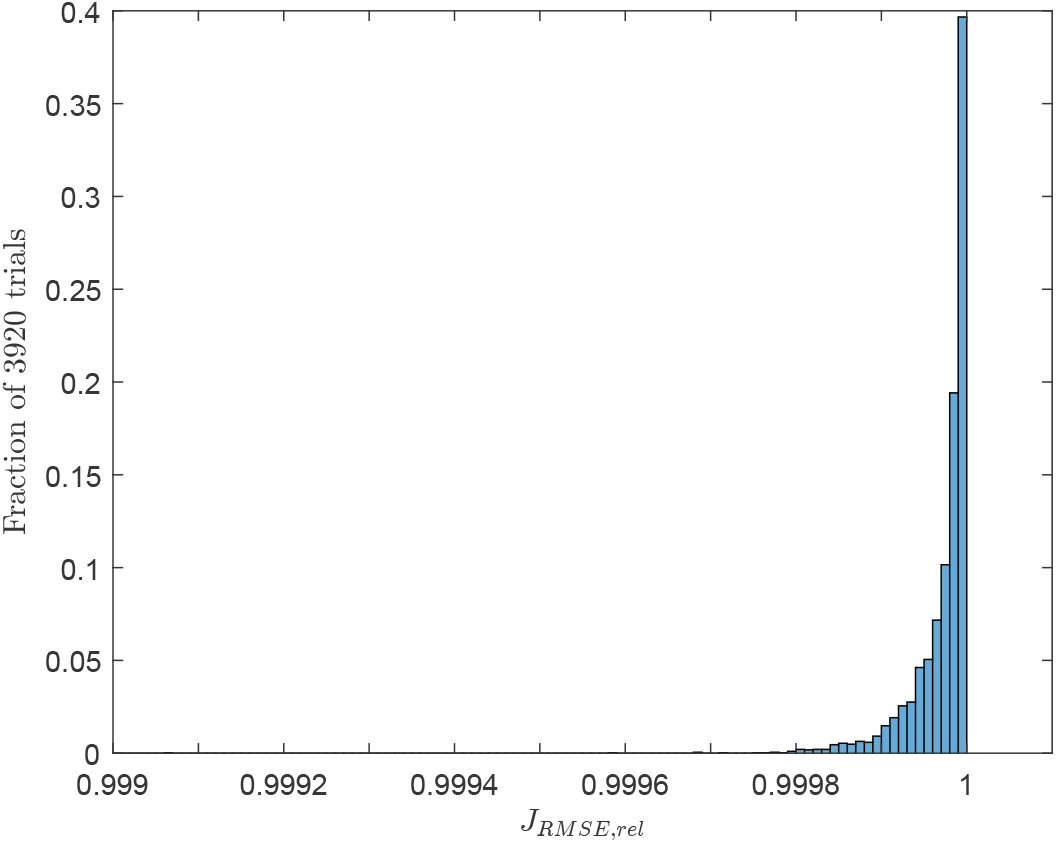
The fractional contribution of the ballistic phase to the overall performance metric (*J*_*RMSE,rel*_) clusters strongly towards 1. 90 % of simulations have *J*_*RMSE,rel*_ *>* 0.9999, and none are below 0.99.

We next define time to stabilisation (*T*_*S*_) as the first timepoint where the instantaneous muscle strain rate is less than 1% of maximum, and the muscle length is within 0.01*l*_0_ of target. Figure S2 shows the correlation between *T*_*T*_ and *T*_*S*_ to RMSE. There is a strong positive correlation between both time metrics and *J*_*RMSE*_, indicating that, in general, performance improves when time to target and time to stabilisation is lower. Note that all trials reach the target and achieve stabilisation within the allotted 0.4 s.

Figure S3 shows correlations between *T*_*T*_, *T*_*S*_, *J*_*RMSE*_ and mean coactivation through each simulation. Coactivation is strongly negatively correlated with each of these measurements. This indicates that, counterintuitively, trials that were fast and stabilized rapidly tended to exhibit higher levels of coactivation overall than trials that were relatively slow.

**Figure S2:**
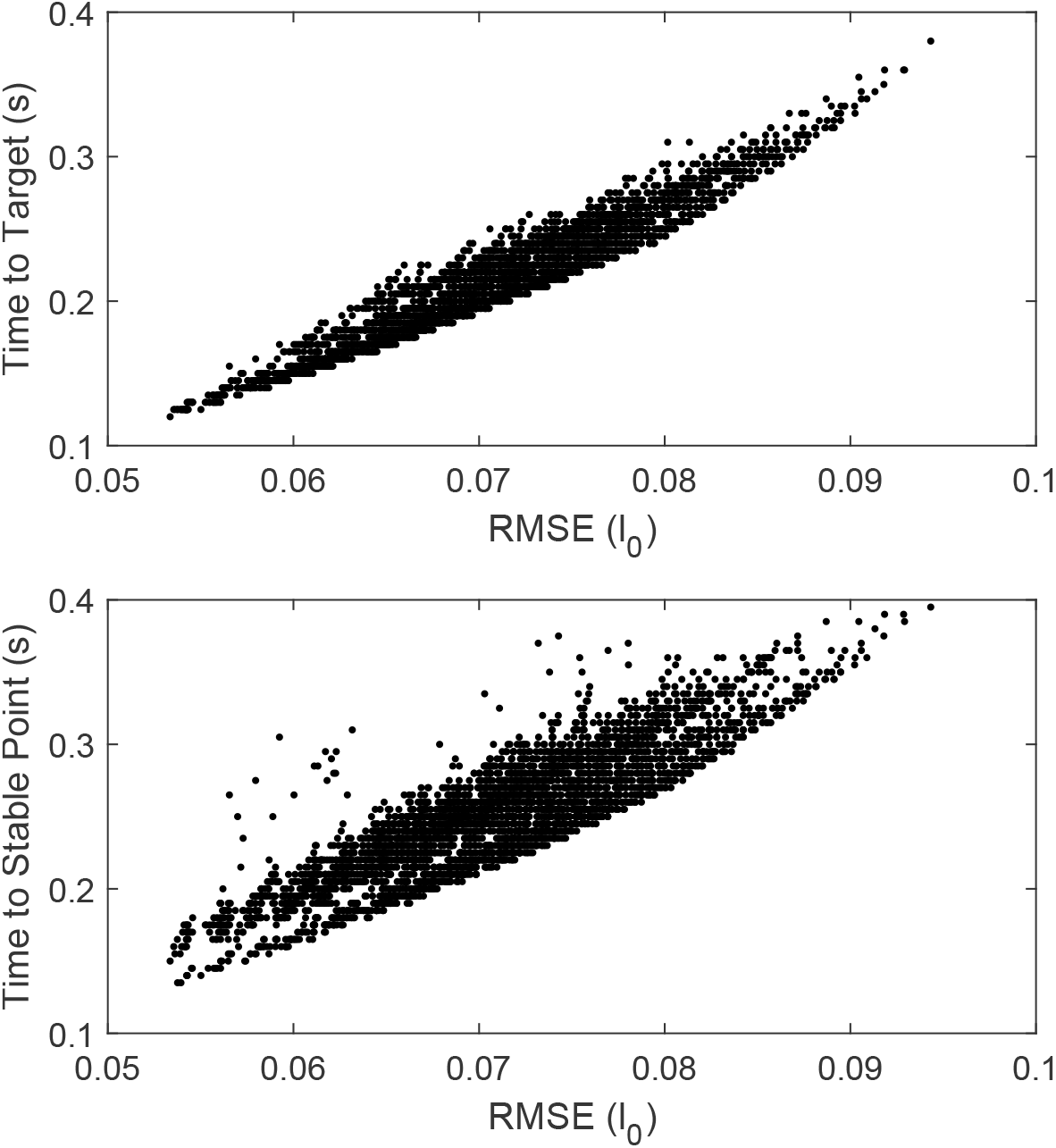
The performance metric of root-mean-square-error (RMSE) is positively correlated with both Time to Target (*T*_*T*_) and Time to Stable Point (*T*_*S*_). Note that all conditions are able to reach either temporal condition within the simulation time window of 0.4 s.

**Figure S3:**
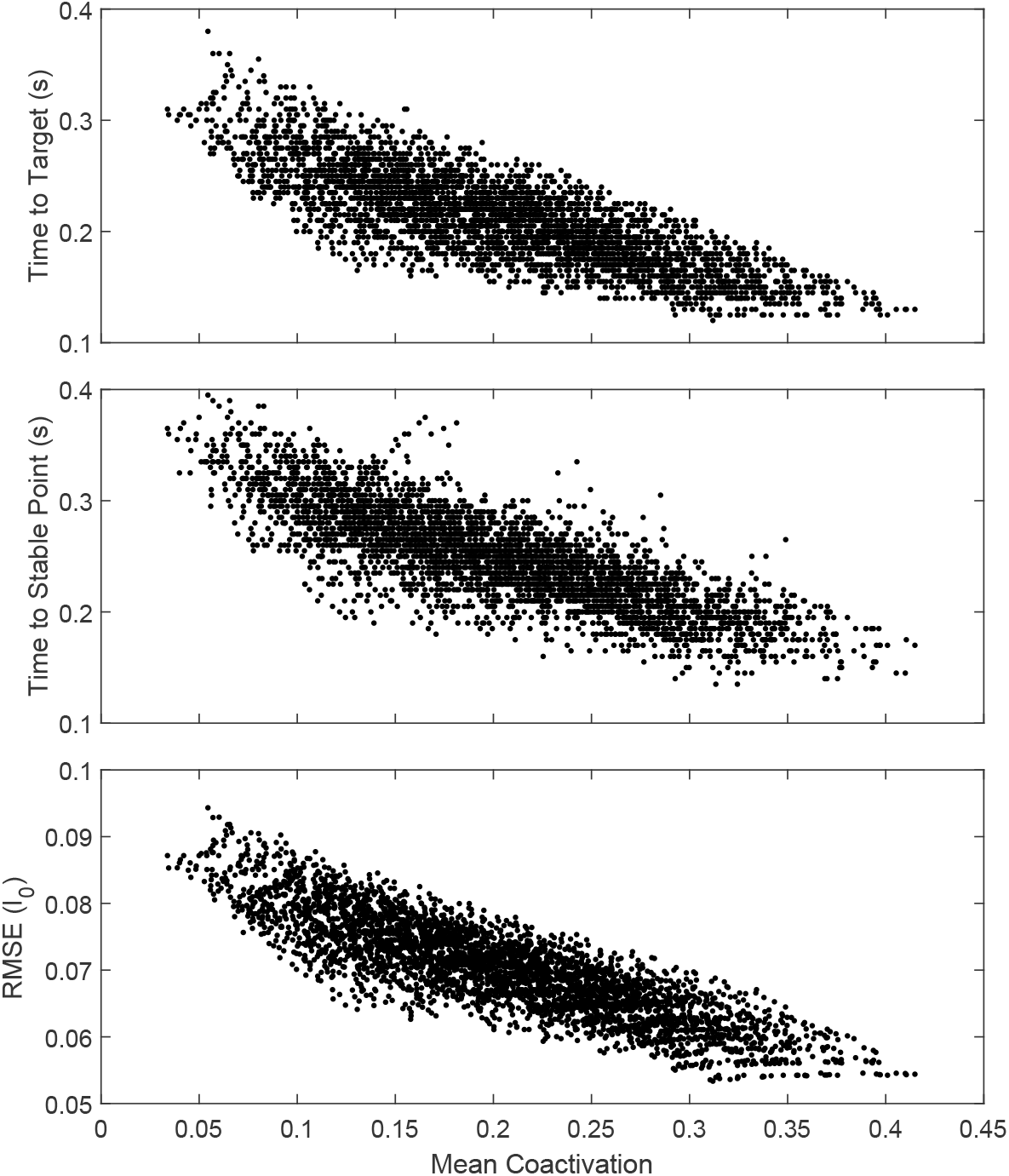
Time to target, time to stable point, and RMSE against mean coactivation for all optimal simulations (3920 conditions). Higher coactivation is associated with lower time to target, lower time to stable point, and lower overall error. By any of these metrics, simulations with higher performance tend to be associated with higher levels of coactivation overall.

